# Treading lightly: Quantitative estimates of seafloor contact for longline trap and hook fishing gear

**DOI:** 10.1101/2024.11.04.621693

**Authors:** Beau Doherty, Lisa Lacko, Allen R Kronlund, Kenyon Alexander, Sean P. Cox

## Abstract

Despite increasing calls for sustainability and ecosystem objectives to manage fishing gear interactions with bottom habitats there are few quantitative approaches for assessing risks from bottom contact fishing. Risk assessments for bottom longline fisheries are particularly challenging due to a lack of information for estimating bottom contact areas from longline gear. In this paper, we demonstrate how data sensors and video cameras deployed on fishing gear can be used to quantify the bottom contact area for longline trap and hook fishing gear from the British Columbia Sablefish fishery. Our bottom contact estimates indicate that Sablefish fishing risks to bottom habitat are low in the majority of fishing areas, since 91.8% of the area fished is expected to have had zero bottom contact over the last 17 years. For the other 8.2% of Sablefish fishing areas that experience some contact from fishing gear, the majority are only contacted once. This indicates that most habitats contacted by Sablefish gear can be expected to have a minimum of 17 years to recover between subsequent bottom contact events. We demonstrate an approach for estimating fisheries bottom contact that can be widely implemented across longline fisheries. Our findings address key data gaps in bottom impacts research for longline gear fisheries, allowing fishing risks to be quantified over fine spatial scales. Such quantitative approaches for habitat risk assessment can provide essential information for management decisions aimed at determining acceptable trade-offs between habitat preservation and fishery benefits.

## 1. Introduction

Vulnerable Marine Ecosystems (VMEs) that contain sensitive benthic habitats such as corals and sponges are increasingly discovered in the world’s oceans (Pham et al., 2014), many in areas that support valuable fisheries. Deep-water corals and sponges provide complex habitat for diverse fish and invertebrates, often within regions of high biodiversity that are sensitive to disturbance (Krieger and Wing, 2002; Heifetz et al., 2009; Buhl-Mortensen et al., 2015). Some species are long-lived, with age estimates of over a century (Andrews et al., 2002, 2009), which makes them susceptible to damage from fishing gear since potential recovery times can span decades. Bottom contact fisheries such as trawl, dredges, and longline gear are perceived as one of the biggest threats to VMEs; however, the potential for bottom fishing impacts on benthic habitats are not well understood. This is particularly the case for longline trap or hook fishing gear known as ‘fixed’ gear that is intended to remain stationary on the seafloor, for which there are limited information to quantify the area of the seafloor that is contacted when gear is deployed and retrieved. If fisheries cannot quantify the damage or risks posed by their fishing gear on seafloor habitats, then they risk losing access to valuable fishing grounds via spatial closures intended to protect VMEs (Barnett et al., 2017). This provides an incentive for fisheries to invest in data collection and science aimed at developing quantitative risk assessments for bottom contact fisheries on VMEs. These bottom contact research efforts can guide management decisions to improve conservation outcomes, while still allowing fisheries to operate in areas where risks are acceptably low.

Bottom impacts from trawl or dredge fisheries are thought to pose a greater risk to VMEs than fixed gears, because they include large nets that are intentionally dragged along the bottom for long distances. While this is likely true when comparing a single trawl or dredging event with that from fixed gear, the total area swept by cumulative longline fishing sets over multiple years can potentially be large depending on the fishing grounds and effort (Welsford and Kilpatrick, 2008; Welsford et al., 2014). Estimates of bottom contact area (i.e., footprints) and damage to benthic communities are more common for trawl or dredges, where it is well established that these gears can remove large quantities of sessile epifauna such as corals, sponges, or sea whips (Sainsbury et al., 1997; Collie et al., 2000; Kaiser et al., 2006b; Clark et al., 2015). In comparison, there are far fewer studies assessing the impact of longline trap or hook fisheries on the seafloor (Eno et al., 2001; Stone, 2006; Stone et al., 2014; Schweitzer et al., 2018), many of which operate in different areas and depths than trawl or dredge fisheries with potential interactions with different species and habitat types (Stone, 2006; Welsford et al., 2014). Fewer still, are studies that attempt to quantify the bottom contact area for longline hook (Ewing et al., 2014; Welsford et al., 2014) and trap (Doherty et al., 2018; Schweitzer et al., 2018) gear, which is necessary for assessing fishery impacts. While bottom trawl or dredge footprints can be quantified as the distance towed multiplied by the width of the trawl gear in contact with the seafloor (Gerritsen et al., 2013; Ewing et al., 2014) estimating bottom contact for fixed gear is more challenging since the extent to which longlines, hooks, and traps move along the bottom during deployment, fishing, and retrieval are poorly understood (Doherty et al., 2018; Stevens, 2021). Longline gear typically includes two end anchors and a longer groundline to which a series of traps or hooks are attached. All of these components have potential to damage sensitive benthic habitats in different ways. For example, traps or anchors can cause damage by landing directly on habitat during deployment or if moved along the seafloor during retrieval (Eno et al., 2001; Stone, 2006; Stone et al., 2014). Epifauna, particularly species with larger and more complex structures (Sampaio et al., 2012), can be hooked or entangled in lines. Finally, groundlines can create shearing forces while moving laterally up to 30 m, significantly increasing the footprint area (Ewing et al., 2014). The extent to which the components of fixed gear move and their potential to damage habitats may vary widely depending on retrieval conditions such as depth, weather, terrain, and currents (Eno et al., 2001; Stone, 2006). All of these factors make it more complicated to calculate the bottom contact area for multiple components of longline trap or hook fishing gear. Given that longline fisheries operate over large spatial scales (Doherty et al., 2018), which may include VMEs or different fishing grounds than trawl or dredge fisheries, it is important to develop robust approaches for quantifying their footprints.

Despite increasing calls for sustainability and ecosystem objectives to manage fishing gear interactions with bottom habitats there are few quantitative approaches for assessing risks from bottom contact fishing. This is largely due to a lack of information on the location of sensitive benthic habitats (SBHs) and the potential for long-term damage caused from different bottom contact fishing gears. The latter is the focus of this paper, for which the first step is to quantify the bottom contact area from fishing gear. We demonstrate how data sensors and video cameras deployed on fishing gear can be used to quantify the bottom contact area for longline trap and hook fishing gear from the British Columbia (BC) Sablefish fishery.

Since 2013, the Canadian Sablefish Association (CSA) has funded bottom contact research that, in collaboration with Department of Fisheries and Oceans Canada (DFO), deploys deep-water cameras, accelerometers, and depth sensors on longline trap Sablefish gear in British Columbia (Fig. 1). The equipment is used to collect observations of deep-sea habitats and to monitor the movement of bottom contact trap gear during fishing operations. The CSA bottom contact research program has revealed four key characteristics of gear movement that are important for understanding longline trap gear interactions with seafloor habitats. First, video observations show that the polypropylene groundlines float and do not contact the bottom, and therefore the traps are the main gear component with potential for bottom contact (Doherty et al., 2018). Second, Sablefish traps are stationary once the gear has settled on the seafloor, during which time their footprint is limited to the bottom area of the trap, but may be less if the trap is on its side. Third, the greatest potential for bottom impacts occurs on gear retrieval, typically the last 1-2 hours of the set, during which traps may move along the bottom (Doherty et al., 2018). The amount of the time the traps drag along the bottom is highly variable across sets ranging from 1-14 min for 25-trap survey sets, based on estimates from a movement classification algorithm using the sensor and video data (Doherty et al., 2018). Finally, conical Sablefish traps have a circular base that is dragged along it’s side during retrieval, limiting the footprint in comparison to rectangular traps of the same width.

**Figure 1:**
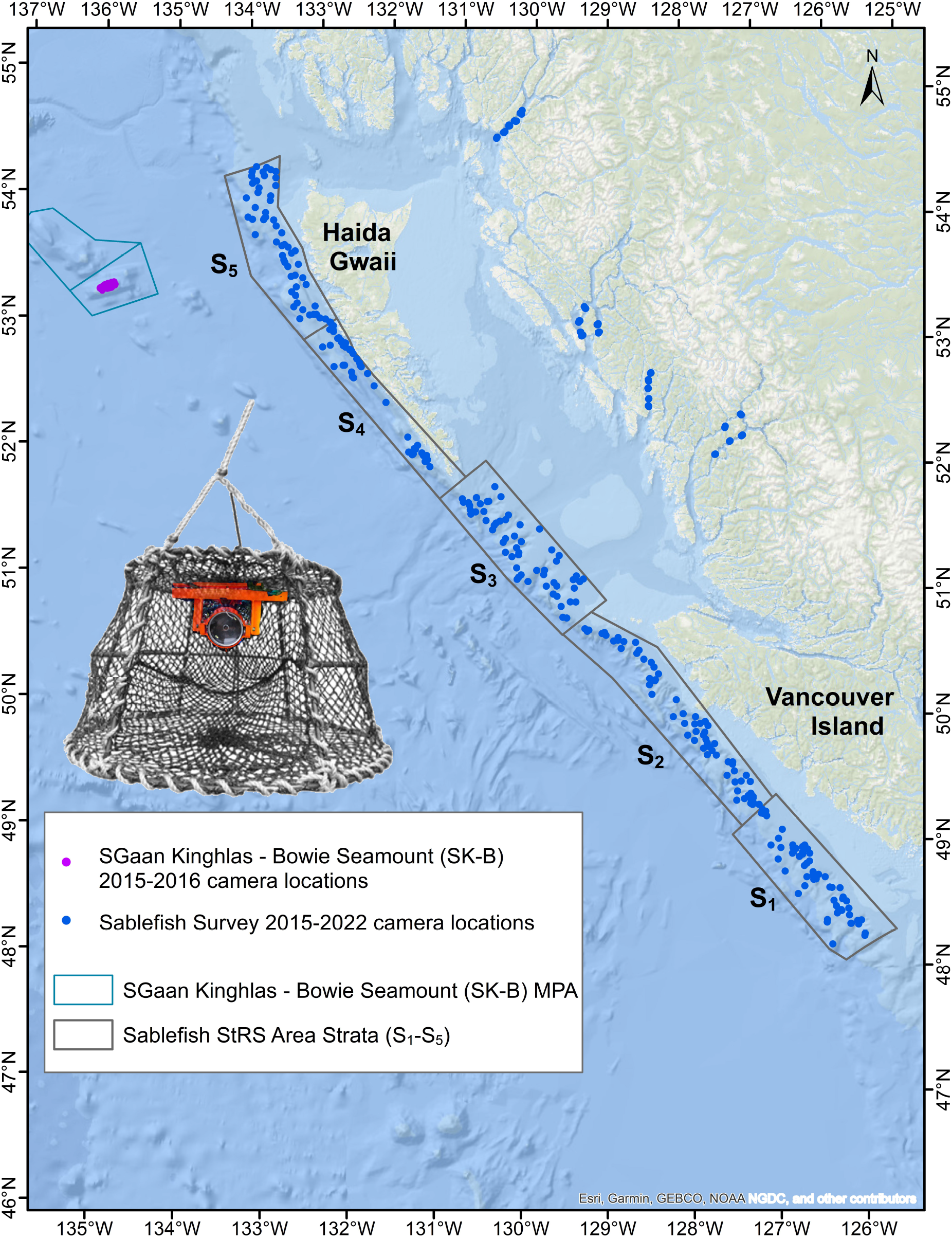
Locations of trap deployments with cameras, accelerometers, and temperature-pressure sensors used in movement classification algorithm. A picture of a mounted camera in a Sablefish trap is shown above the legend on the left side of the map.

Findings to date can be used to develop and demonstrate an adaptive management approach to improving sustainable fishing practices for bottom contact fisheries. This includes i) an integrated system of data capture technology for sensitive benthic habitats, ii) in-situ monitoring and estimation methods to quantify bottom contact from fishing gear, iii) presence-absence modelling to map SBHs, iv) risk assessment for quantifying bottom fishing impacts on SBHs and spatial assessment of conservation status, and v) a simulation framework for evaluating fishing impacts on SBHs and their recovery trajectories under alternative management approaches. This paper focuses on i) and ii), demonstrating a quantitative approach to estimating bottom contact for fixed gear that can be widely applied to any type of longline trap or hook fishing gear. To achieve this, we updated the classification algorithm for estimating trap movement from (Doherty et al., 2018) and applied it to a larger dataset that includes observations from commercial and survey fishing trips from 2015-2022. We assessed the accuracy of the movement classification algorithm by comparing the estimated time that the trap moves along the seafloor with the true time captured from continuous video observations on select sets. Additionally, we evaluated whether different fishing conditions such as depth or the trap position along the groundline effects gear bottom contact. This information was then used to estimate the total bottom contact area for the BC Sablefish fishery over 17 years from 2007-2023. These estimates provide essential information for risk assessment and management decisions in BC for fine spatial scales. We demonstrate an approach for data collection and footprint estimation that can be widely applied to other longline fisheries.

## 2. Methods

### 2.1. Data collection

We deployed accelerometers (Actilife wGT3x-BT monitors), temperature-pressure recorders (Sea-Bird SBE 39), and a deep-water camera system (Doherty et al., 2018, 2021) on traps deployed on longlines fished during the annual stratified random survey (StRS) for BC Sablefish from 2015-2022, as well as during commercial fishing trips to the SGaan Kinghlas - Bowie Seamount (SK-B). The longline trap gear used on the survey includes a 1200 m longline (Wyeth et al., 2007) with 25 or 26 traps (26 traps for sets with cameras), whereas commercial gear typically consists of 60 traps on longlines around 3 km long (Doherty et al., 2018). The accelerometers, depth sensors, and cameras were used to collect information on bottom substrate and gear movement during deployment, soak, and retrieval of selected sets. The deep-water ‘trap camera’ system is composed of a GoPro HD Hero, four LED lights, and a lithium-ion battery pack, depth sensor, tri-axial accelerometer, circuit board, and controller, all of which are contained in a 3.6 kg stainless steel housing that can be deployed as deep as 1500 m (Doherty et al., 2018).

Cameras deployed from 2015 to 2020 were programmed to record 1-minute videos at 2 hour intervals or when accelerometers within the camera housing triggered video recordings during gear movement at impact forces greater than 0.6 g (Doherty et al., 2018). These camera settings were intended to conserve battery power throughout the typical 24-hour set duration of the survey, enabling the collection of in-situ observations of gear and habitat throughout the set.

Camera settings were adjusted for StRS gear deployments from 2021 to 2022 to focus on video observations during gear retrieval, particularly, when there is potential for traps to move along the seafloor. Video recordings of up to 2000 seconds were triggered by the accelerometer motion-activation threshold (0.6 g), which allowed for continuous video from the start of gear retrieval to the moment when the trap containing the camera was hauled out of the water. These revised settings also included a 21.5 hour time delay to conserve battery power and limit any video recording prior to gear retrieval. Video clips were reviewed to classify observations of gear behaviour as either stationary, dragging, or suspended. The 2021-2022 survey protocols were also modified to add experimental sets with 60-traps for monitoring gear bottom contact, in addition to the typical 25-trap survey sets.

### 2.2. Movement classification algorithm

We developed a movement classification algorithm that uses depth sensor and accelerometer data to classify different types of trap movement during gear retrieval (Doherty et al., 2018). Processed video data are also used to confirm the movement classification, when available, but are not essential for applying the algorithm. The movement classification algorithm is applied to data collected during the last two hours of fishing. The approach originally described in (Doherty et al., 2018) was updated for classifying gear behaviour during one second intervals over the last two hours (7200 seconds) of each fishing set using the following measurements:

- depth sensor measurements (*d*_*t*_),
- depth change per second (Δ*d*_*t*_),
- depth change over 10 s intervals (*d*_*t*_ − *d*_*t*−10_),
- acceleration (g) at 1 s intervals (*a*_*t*_),
- acceleration variance over 10 s intervals using 10 Hz acceleration 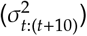,
- and observed trap movement from 1-min video intervals (*V*_*t*_).

Depth sensor measurements were available for 1, 3, 5, or 10 second intervals depending on the set.

Trap movement was classified as either “stationary”, “dragging”, or “suspended” (Fig. 2), using the following algorithm:

**Figure 2:**
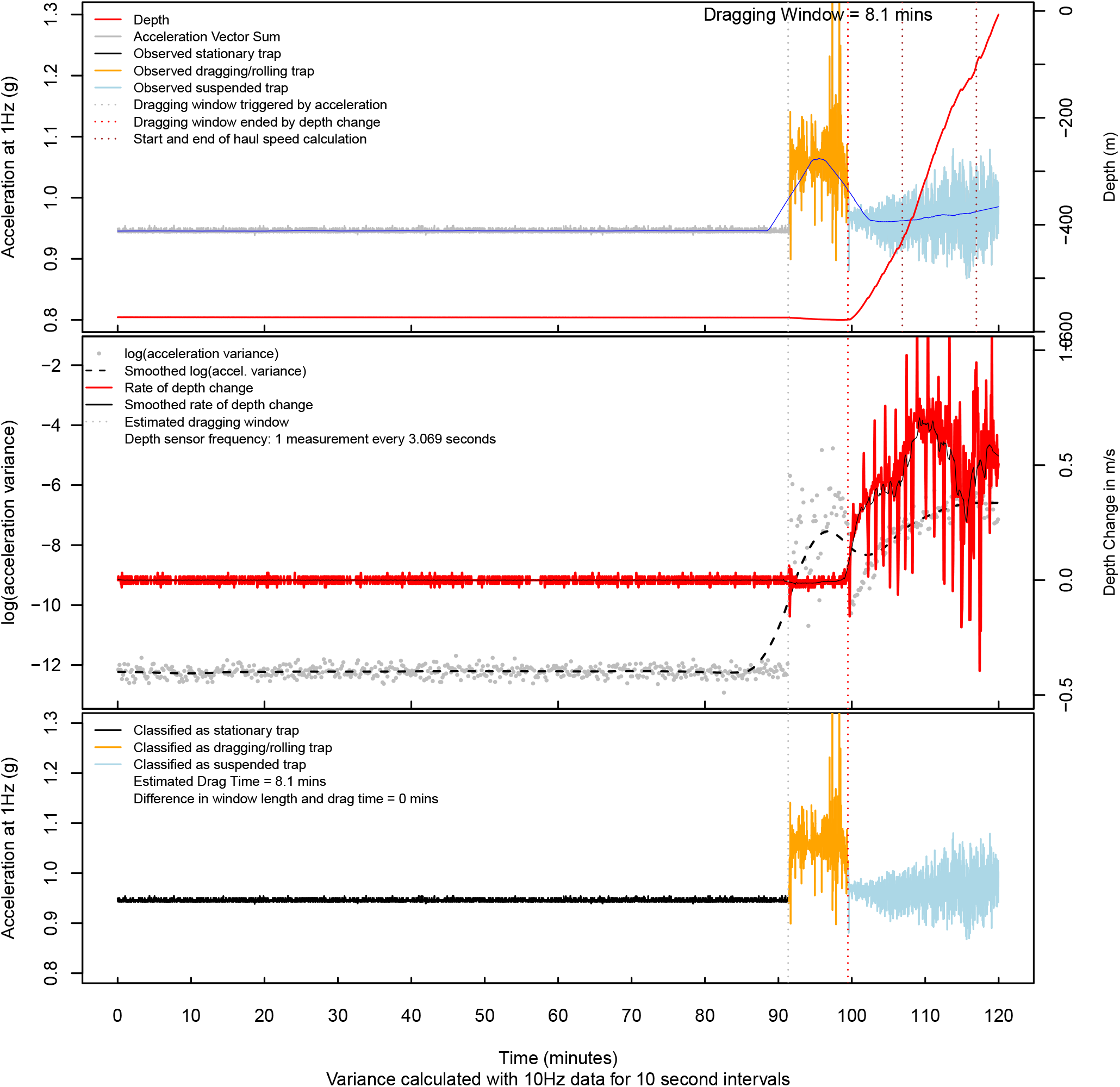
Example of movement classification algorithm inputs (top, middle) and outputs (bottom) for a trap deployment on the 2022 survey where the estimated drag time is the same as the observed drag time from video. Continuous video observations of trap movements are also shown in top panel, which for 2021-2022 were held out from the classification algorithm so that they could be used for model validation.

**Step 1**. Define start and end times for the period where the trap has potential to drag along the seafloor during gear retrieval (i.e., ‘drag window’):

a. Choose the start of the drag window *t*^start^ as the first time step subject to (s.t.) any one of three conditions being true

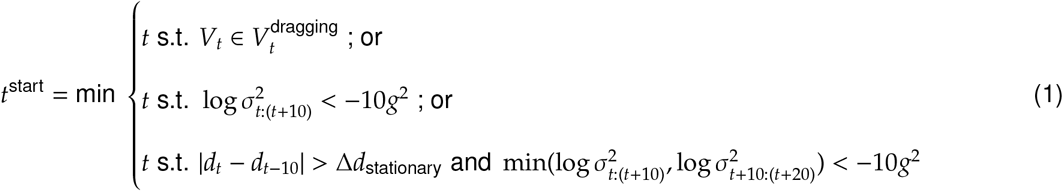

where 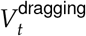 represents the first time-step of video where the trap is dragging. The Δ*d*_stationary_ term is the maximum change in consecutive depth measurements while the trap is stationary (typically < 5 cm/s), which can trigger the drag window up to 20 seconds prior to the 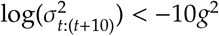 condition.
b. Choose the end of the drag window *t*^end^ as the first time-step subject to any one of three conditions being true

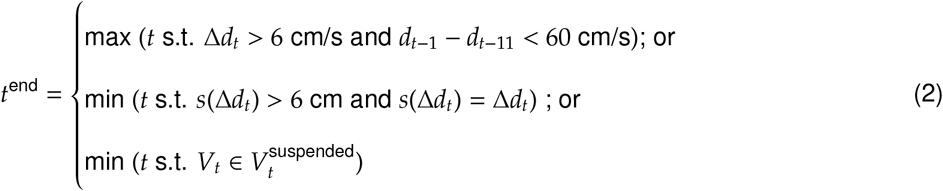

The 1st condition indicates the latest time step where depth change > 6 cm/s and where the mean depth change is < 6 cm/s for the preceding 10 seconds. The 2nd condition indicates the first time-step where the 30-second moving average for depth change *s*(Δ*d*_*t*_) exceeds 6 cm/s and intersects with the depth change per second. Finally, the 3rd condition is the first time-step where there is video of the trap suspended in the water column 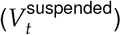.

**Step 2**. Determine the class (*C*_*t*_) of the trap behaviour for each interval between depth sensor measurements (Δ*t*^∗^) within the drag window, where 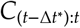 intervals are classified as:

i. “stationary” when 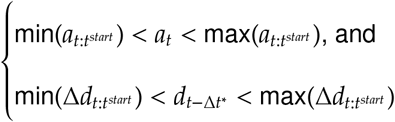 ;
ii. “suspended” when 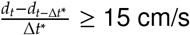 and
iii. “dragging” when *C*_*t*_ ≠ ‘stationary’ and *C*_*t*_ ≠ ‘suspended’

where Δ*t*^∗^ is 1, 2, 3, or 10 seconds depending on set. The specific thresholds for depth change and acceleration variance used in step 1 and step 2 [eqns (1)-(2)] were determined iteratively by fine-tuning the algorithm based on visual inspection of performance from time series plots of the movement classification inputs and outpus (Fig. 2)

**Step 3**. Set the classes of trap behaviour to 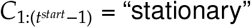 for each *t* prior to the drag window and 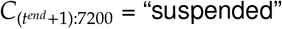 for each *t* after the drag window.

Video clips of dragging and suspended traps provided lower and upper bounds for the drag window. The difference between the start and end times of the drag window is considered the maximum potential dragging time, which is equal to estimated drag time in cases where neither conditions i) or ii) from strep 2 are triggered.

The distance that a trap is dragged along the bottom *l* was estimated by multiplying the estimated drag time by the estimated hauling speed for each set, which we used to approximate the speed at which traps drag along the bottom. Haul speed (*v*) was calculated as the median rate of depth change (cm/s) during gear retrieval between depths at 75% of the bottom depth 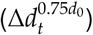 and 100 m 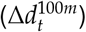 below the surface:

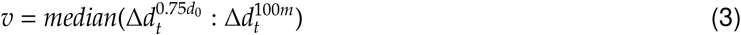

To assess algorithm performance, we compared estimated drag times from the movement classification algorithm with the true drag time observed from continuous video of gear retrieval from *K* = 19 traps deployed on 2021-2022 StRS sets.

We calculate the relative error *RE*_*k*_ for each *k* = 1, 2, …, *K* trap as:

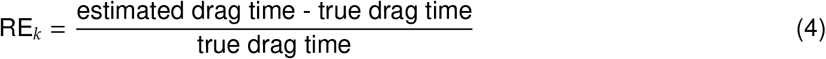

We then calculate the mean relative error 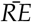, weighted by the true drag time *w*_*k*_ as

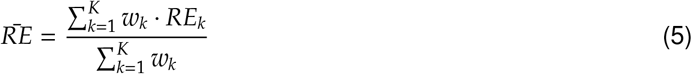

### 2.3. Fishery bottom footprints

Here we described the necessary steps to estimate the bottom contact area for Sablefish longline trap and longline hook fishing gear. This involves developing an approach for estimating footprints for i) longline trap sets, ii) longline hook sets, iii) and an overall approach for calculating the combined footprints of cumulative sets from both gear types over multiple years, that considers potential for sets to have overlapping contact areas.

The commercial fishery data used for the footprint analysis consisted of trips from the Sablefish sector, as well as combined trips for Pacific Halibut and Sablefish sectors sourced from the fisher logbooks stored in the Fishery Operations System (FOS) enterprise database of the DFO. The data presented in this report adhere to the regulations outlined in the Access to Information and Privacy Act. Figures are presented in a format complying to these regulations for the time period and grid cell area of interest, whereby grid locations with fewer than three unique fishing vessels are not shown (i.e., ‘the rule of three’, Tomasic, 2023). For simplicity, we refer to these data throughout the report as Sablefish longline trap and hook fisheries, even though the longline hook component contains both directed Sablefish and combined Pacific Halibut/Sablefish trips.

We estimated the coastwide bottom footprint for longline trap and longline hook gear from the BC Sablefish fishery using two different methods. The first method assumes no-overlap of bottom contact area within a 4×4 km grid cell, such that the total bottom contact area for each grid cells is the sum of the footprint *f* for all sets (*j* = 1, 2, …, *J*) of each gear type *g* in grid cell *i*

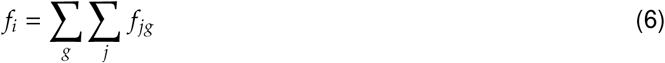

The proportion of the grid cell *λ*_*i*_ that is contacted by fishing gear is the footprint divided by the total grid cell Area *A*_*i*_, which is 16 km^2^ for a 4 × 4 km grid cell.

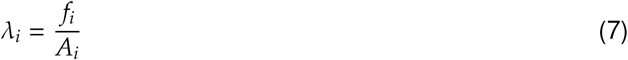

If *f*_*i*_ > *A*_*i*_, then *λ*_*i*_ > 1 and there must be some overlap of set footprints within the cell, in which case the no-overlap approach sets *f*_*i*_ equal to *A*_*i*_.

The second ‘overlap’ method assumes that sets are randomly distributed in each grid cell according to a Poisson distribution (Gerritsen et al., 2013; Amoroso et al., 2018) and that there is some overlap of bottom contact from multiple sets within a grid cell. The proportion *P* of a grid cell that is contacted *h* times (Gerritsen et al., 2013) is defined as :

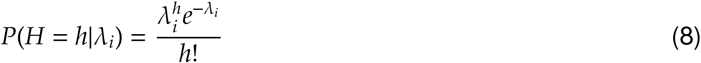

The proportion of a cell that is contacted at least once (i.e., *h* ≥ 1) is then:

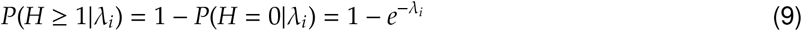

Therefore, to account for random overlap of set footprints within a grid cell, we modify eqn. (6) to estimate the footprint areas contacted at least once for each *i* cell as:

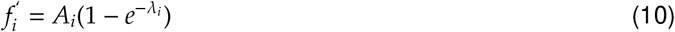

We also calculate the proportion of grid cells contacted at least two or three times via a cumulative distribution function for a gamma distribution calculated using the pgamma function in R (S2 in Gerritsen et al., 2013):

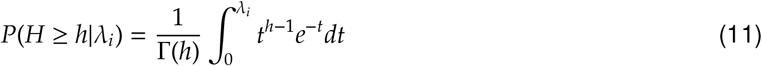

We use the overlap approach to generate maps of bottom contact for the BC Sablefish fishery for longline trap and longline hook gear deployed from 2007-2023. We included fishing trips for i) Sablefish longline trap or hook fishing (K licenses), and ii) Pacific Halibut/Sablefish combination trips (K/L licenses) using longline hook gear.

We calculate aggregate coastwide footprint for the fishery by summing bottom contact area within each grid cell *i*

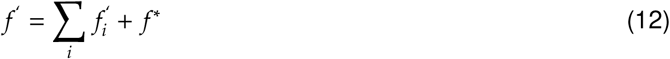

where *f* ^∗^ represents the bottom contact area for all sets with unknown grid locations, which are calculated using the no overlap approach in eqn. (6). This includes sets lacking deployment location data, as well as set locations that were withheld for privacy reasons. The latter applies to any grid locations that did not have set deployments from at least three unique fishing vessels.

#### 2.3.1. Sablefish longline trap gear contact

The bottom contact area for a typical Sablefish conical trap with a 137 cm (54-inch) bottom hoop diameter can be calculated by multiplying the footprint length *l*_*n*_ for trap *n* on set *j*, described in subsequent section, by the footprint width *w*

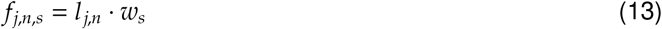

where *w*_*s*_ are taxa-specific *s* footprint widths for contact that depend on the morphology of habitat forming species (described in trap footprint widths below). The *n* indicates the trap retrieval number for a set with n=1,2,..,*N* traps, for which *N*=60 total traps for a typical commercial Sablefish set and *N*=25 for survey sets.

We then calculate the bottom contact *f*_*j,s*_ for a longline trap set *j* and taxa *s* as the sum of each trap footprint on the set

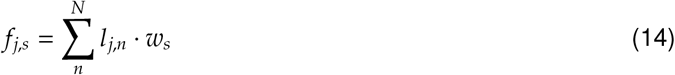

Details for calculating the trap footprint length *l* and footprint widths *w*_*s*_ are described below. Footprint lengths are estimated via linear models fit to outputs from the movement classification algorithm that account for trap position along the groundline and depth effects. Taxa-specfic footprint width *w*_*s*_ scenarios are based on the probability of trap-contact with different benthic habitats that depend on their height distribution.

##### 2.3.1.1. Trap position and depth effects on footprint lengths

We fit a linear model to estimate the effects of trap position on the longline set (e.g., start, middle, end) and set depth on estimates of trap footprint lengths *l*_*j,n*_ from the movement classification algorithm for trap *n* and set *j*:

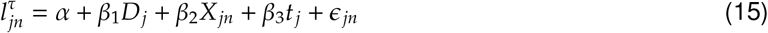

where *α* is the intercept term and *β* terms are estimated coefficients for predictor varables for depth *D*_*j*_, trap retrieval position *X*_*n*_, and survey year *t*_*j*_. The τ variable is used to transform the trap footprint length estimates from the classification algorithm so that they approximate a normal distribution (Tukey Ladder transformation, τ = 0.3, Shapiro-Wilk *p* = 0.45). Trap drag length estimates, depth, and trap position for the same sets were averaged for model fitting, so that the model only includes one observation per set. We excluded one outlier set from model fitting that was confirmed as a misclassification via video (discussed in results).

The trap position on retrieval *n* was calculated relative to the total number of traps on the set *N* and expressed as a percentage of set length:

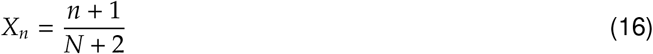

We add 1 to the numerator and 2 to the denominator to account for spacing between anchors at both ends of the set. SK-B trips were excluded from this analysis because these data don’t include trap numbers on retrieval.

We use the linear model fit from eqn. 15 to estimate the footprint lengths of longline trap gear deployed by the BC Sablefish fishery from 2007-2023, for an average year effect. For 6% of longline trap sets where the depth was unavailable but deployment locations were geo-referenced, an estimated depth was extracted from the General Bathymetric Chart of the Oceans (GEBCO_2014 Grid, version 20150318, www.gebco.net) at the set midpoint. For 0.03% of sets where both deployment locations and depth were unknown, we used the median depth of 604 m for longline trap sets from 2007-2023. When the number of traps deployed on a set were unavailable, we assumed 60 traps per longline set.

The total footprint length for the longline trap set *l*_*j*_ is then the sum of *n* = 1, 2, …, *N* trap footprint lengths on the set:

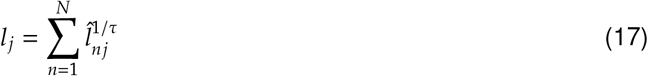

##### 2.3.1.2. Trap footprint widths

The potential for trap contact with habitat forming species is dependent on two factors: i) the width of the furrow created by the dragging trap, and ii) whether the edges of the trap outside the furrow are higher than the height of the organism (Fig. 3). For organisms inside the furrow path there is 100% probability of contact, while organisms outside the furrow path have a decreasing probability of contact *ρ* for increasing distances from the centre of the trap and decreasing height of sessile organisms. We estimate the expected trap-coral *w* contact width for 5 mm intervals *i* = −685, −680, …, 685 from the centre of a 1.37 m trap at *x* = 0:

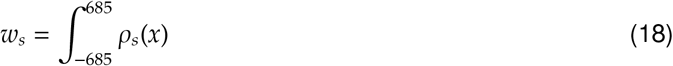

where *ρ*_*s*_(*x*) is the proportion of the population of sessile organisms *s* with heights greater than the height of the trap off-bottom at interval *x*.

**Figure 3:**
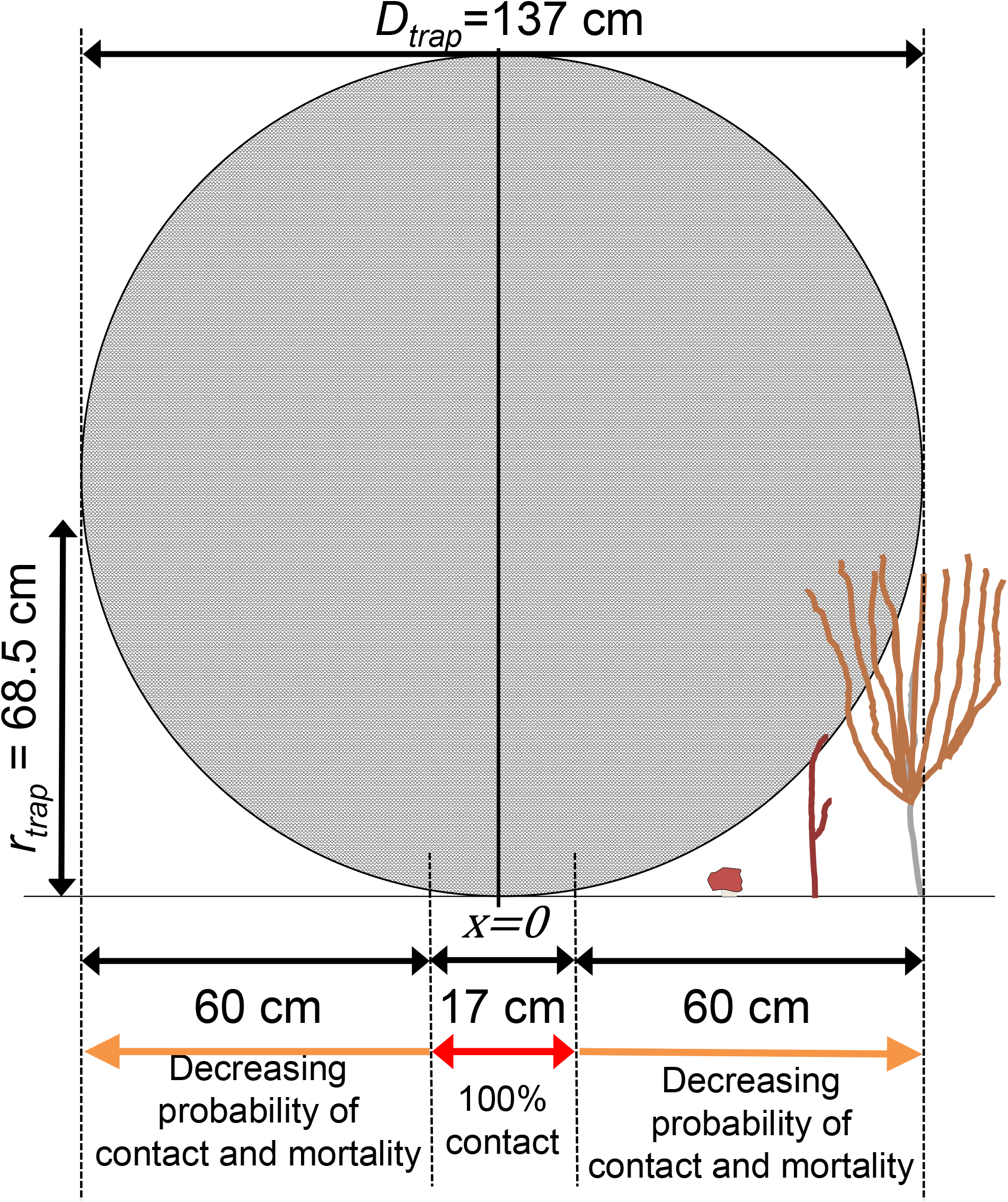
The view of the underside of an overturned Sablefish trap (1.37 m diameter) on the seafloor while dragging during gear retrieval and the potential trap-coral contact width outside the furrow path (17 cm) for small (h = 6 cm), medium (h = 28 cm), and large (h = 48 cm) coral taxa. The probability of trap-coral contact and the level of damage decreases with distance from the centre of the trap and decreasing height of corals.

We use furrow width estimates of 17 cm (95% CI=15-19) from Doherty et al. (2018) to define the *x* interval with *ρ*_*s*_ = 100%. For *x* intervals outside the furrow width, we calculate *s* taxa-specific *rho*_*s*_*x* scenarios according to observed height distributions (unpublished data, C. Rooper, DFO, Fig. 4) from surveys in Alaska (Rooper et al., 2016; Wilborn et al., 2018; Sigler et al., 2023) for corals (Order Alyconacea), sponges (Phylum Porifera), sea whips (Order Pennatulacea), and hydrocorals (Order Anthoathecata), which are the most commonly observed taxanomic groups from trap camera observations from the BC Sablefish survey (Gemmell, 2022). The Alcyonacea group was further split into 4 specific familes (Isididae, Paragorgiidae, Plexauridae, Primnoidae), for which there was more detailed information on height distribution (Table 2). Finally, we generate a weighted footprint *w* value across all *S* = 7 taxa, according to the mean proportion π_*s*_ of each taxon observed from trap camera observations in BC (Table 2) that was used for calculating the Sablefish fishery footprint.

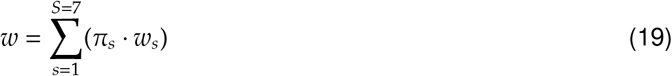

**Table 1:**
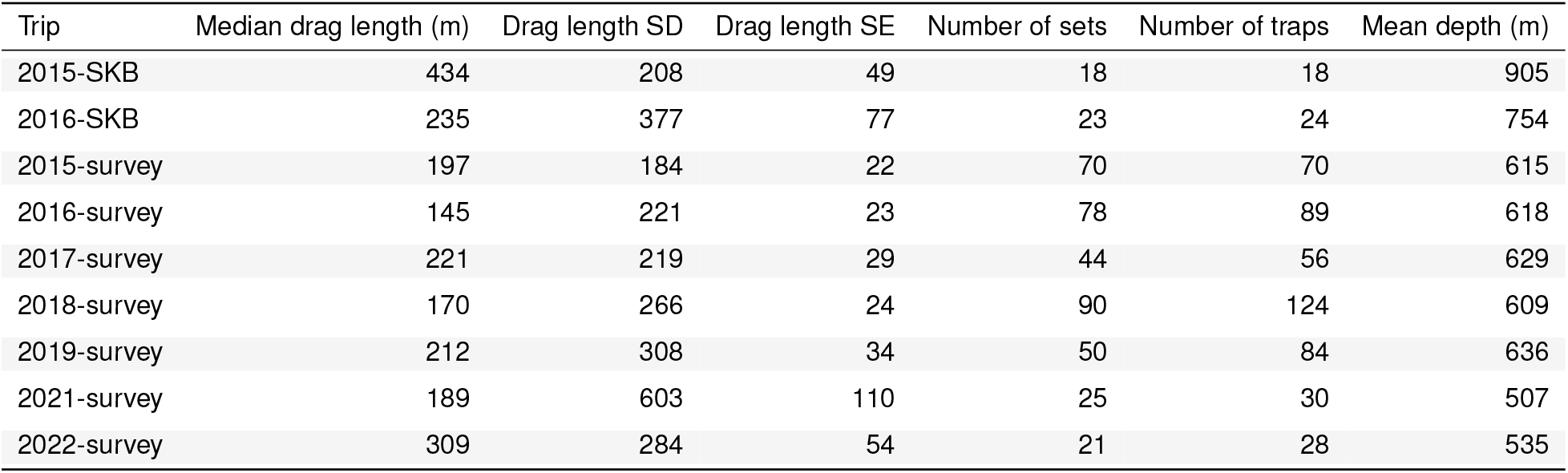
Raw summary statistics for estimated trap drag lengths (m) and mean depths (m) from traps deployed with sensors on Sablefish survey and SK-B fishing trips from 2015-2022. The total number of sets and traps deployed with sensors are shown for each year and trip type.

| Trip | Median drag length (m) | Drag length SD | Drag length SE | Number of sets | Number of traps | Mean depth (m) |
| --- | --- | --- | --- | --- | --- | --- |
| 2015-SKB | 434 | 208 | 49 | 18 | 18 | 905 |
| 2016-SKB | 235 | 377 | 77 | 23 | 24 | 754 |
| 2015-survey | 197 | 184 | 22 | 70 | 70 | 615 |
| 2016-survey | 145 | 221 | 23 | 78 | 89 | 618 |
| 2017-survey | 221 | 219 | 29 | 44 | 56 | 629 |
| 2018-survey | 170 | 266 | 24 | 90 | 124 | 609 |
| 2019-survey | 212 | 308 | 34 | 50 | 84 | 636 |
| 2021-survey | 189 | 603 | 110 | 25 | 30 | 507 |
| 2022-survey | 309 | 284 | 54 | 21 | 28 | 535 |

**Table 2:** Summary statistics for height observations (cm), trap footprint *w*_*s*_ widths (cm), and weightings π_*s*_ (%) of major taxonomic groupings used in weighted footprint. Height observations (unpublished data, C. Rooper, DFO) are from underwater surveys in Aleutian Islands (2012-2014, Wilborn et al. 2018), Eastern Bering Sea (2014, Rooper et al. 2016), and Gulf of Alaska (2013, 2015, 2017, Sigler et al. 2023).

| Type | Taxonomic group | n observations | mean height | SD height | min. height | max. height | $w_s$ (cm) | $\pi_s$ |
| --- | --- | --- | --- | --- | --- | --- | --- | --- |
| Alcyonacea Corals | Isididae | 104 | 35.8 | 19.9 | 9.7 | 116.2 | 113 | 2.8 |
| Alcyonacea Corals | Paragorgiidae | 62 | 35.1 | 24.1 | 4.9 | 116.3 | 108 | 4.7 |
| Alcyonacea Corals | Plexauridae | 822 | 26.4 | 13.3 | 3.5 | 88.2 | 103 | 3.7 |
| Alcyonacea Corals | Primnoidae | 5374 | 28.9 | 15.5 | 2.0 | 127.5 | 106 | 3.7 |
| Sponges | Porifera | 11282 | 22.6 | 13.5 | 1.4 | 288.2 | 96 | 28.3 |
| Sea Whips | Pennatulacea | 4979 | 55.3 | 51.5 | 1.7 | 266.3 | 104 | 44.4 |
| Hydrocorals | Stylasteridae | 1576 | 9.8 | 7.1 | 1.3 | 111.6 | 67 | 12.4 |

**Figure 4:**
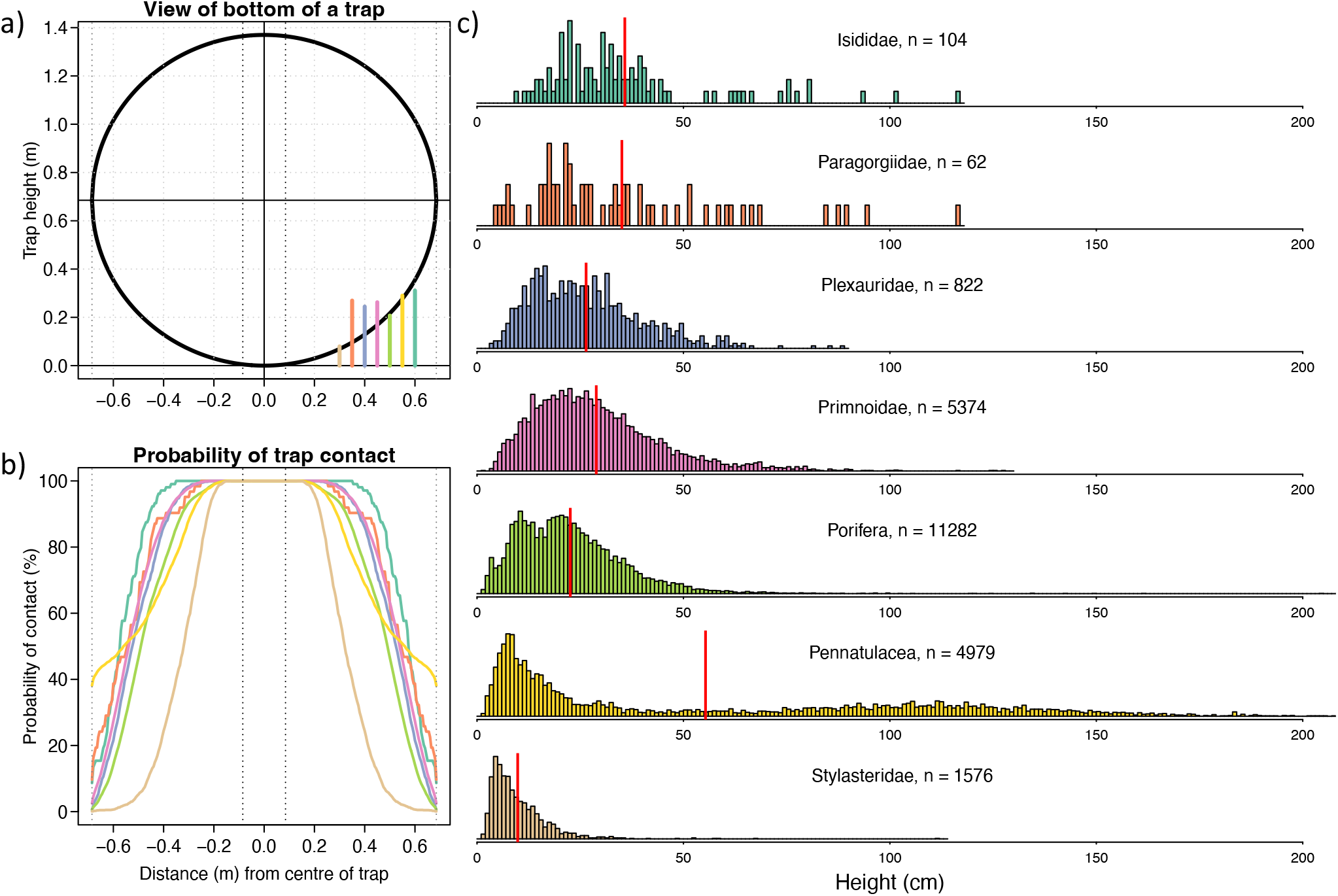
a) View of the underside of an overturned Sablefish trap (1.37 m diameter) on the seafloor during gear retrieval and b) probability of trap contact for different taxa according to height distributions in c). Vertical dashed lines at +/-8.5 cm and +/- 68.5 cm indicate the 17 cm furrow width and trap edges, respectively. Vertical coloured lines indicate median height from taxa distributions. c) Height distributions from Alaska for Alcyonacea coral families (Isididae, Paragorgiidae, Plexauridae, Primnoidae), sponges (Porifera), sea whips (Pennatulacea), and hydrocorals (Stylasteridae). The vertical red line indicates the mean height for each taxon. Height observations (unpublished data, C. Rooper, DFO) are from underwater surveys in Aleutian Islands (2012-2014, Wilborn et al. 2018), Eastern Bering Sea (2014, Rooper et al. 2016), and Gulf of Alaska (2013, 2015, 2017, Sigler et al. 2023).

#### 2.3.2. Sablefish longline hook gear

The bottom contact area of a Sablefish longline hook gear set *j* is estimated as the total set length *l* multiplied by the lateral line movement (*w*) during gear retrieval:

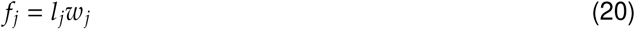

When the set length was unavailable, we use the median set length of 3.1 km for Sablefish longline hook gear from 2007-2023.There are few empirical data for longline gear bottom contact and no specific studies exist for BC fisheries; however, bottom contact estimates of similar longline hook gear are available for Patagonian Toothfish (*Dissostichus eleginoides*) fisheries in Antarctica (Ewing et al., 2014). The bottom contact area of longline gear was estimated as the total set length (median of 8.8 km) multiplied by the average lateral line movement, which was estimated as 6.2 m (SE=1.9) from cameras deployed on longline gear (Ewing et al., 2014; Welsford et al., 2014). For example, using a mean footprint width of 6.2 m and the median set length of 3.1 km in the Sablefish fishery from 2007-2023 generates a footprint estimate of 19,220 m^2^, which is 0.1% of the area within a 4×4 km grid cell. The lateral line movement may be affected by factors such as the line tension, seafloor bathymetry, and the vessel position relative to the lifting point (Ewing et al., 2014). To account for the uncertainty in the lateral line movement estimates within the BC Sablefish fishery, we estimated longline footprints using 3 different scenarios for the lateral line movement *w*, which include a baseline scenario using the mean estimate of 6.2 m from (Ewing et al., 2014) and scenarios corresponding to approximate 95% confidence intervals for the mean, estimated as 6.2 m +/- 1.96 SE.

## 3. Results

### 3.1. Movement classification algorithm

The movement classification algorithm was applied to 523 traps deployed with sensors on commercial fishing sets at SK-B from 2015-2016 and StRS sets from 2015-2022. Summary statistics for estimated drag lengths by year and trip type are provided in Table 1.

Where available, we compared the 2021-2022 estimates of drag time from the classification algorithm with the true value obtained from video for 19 trap deployments (Fig. 6). The majority of the estimates were within 1 minute of the true value; however, there were 5 estimates that were not. These included 3 overestimates between 1.9 and 4 minutes, one underestimate of 4.1 minutes, and one large outlier from the 2021 survey that overestimated drag time by 22.7 minutes. The latter trap was estimated to have a 33.8 minute drag window, all of which was classified as dragging. When logging the video data, more than 20 minutes of the video was classified as stationary with a high degree of confidence since the seafloor was clearly visible with a Sea Whip (Pennatulacea) as a reference object. However, on closer inspection this trap was not completely stationary. It was swaying and moving very slowly in relation to the reference object, moving less than 0.5 m over 20 minutes. In this case, it appears that the swaying and slow movement may have contributed to the algorithm’s significant overestimate, resulting in the classification of the entire drag window as ‘dragging’. Estimates of relative error were very sensitive to this one outlier with a weighted mean relative error that was 3.8% when excluding the outlier and 20% when it was included. Since this outlier was confirmed by video as a misclassification, it was removed from subsequent analyses investigating depth and trip effects on gear movement.

### 3.2. Trap and depth effects on gear movement

Linear model results found evidence that both depth (p<0.001, ANOVA III) and trap retrieval order (p<0.001, ANOVA III) had an effect on trap drag length (Figure 5). The drag time effect indicates that traps drag less on the bottom for deeper depths, while the trap position effect indicates traps drag less when retrieved near the beginning of the set (i.e., closer to the first anchor buoy on retrieval).

**Figure 5:**
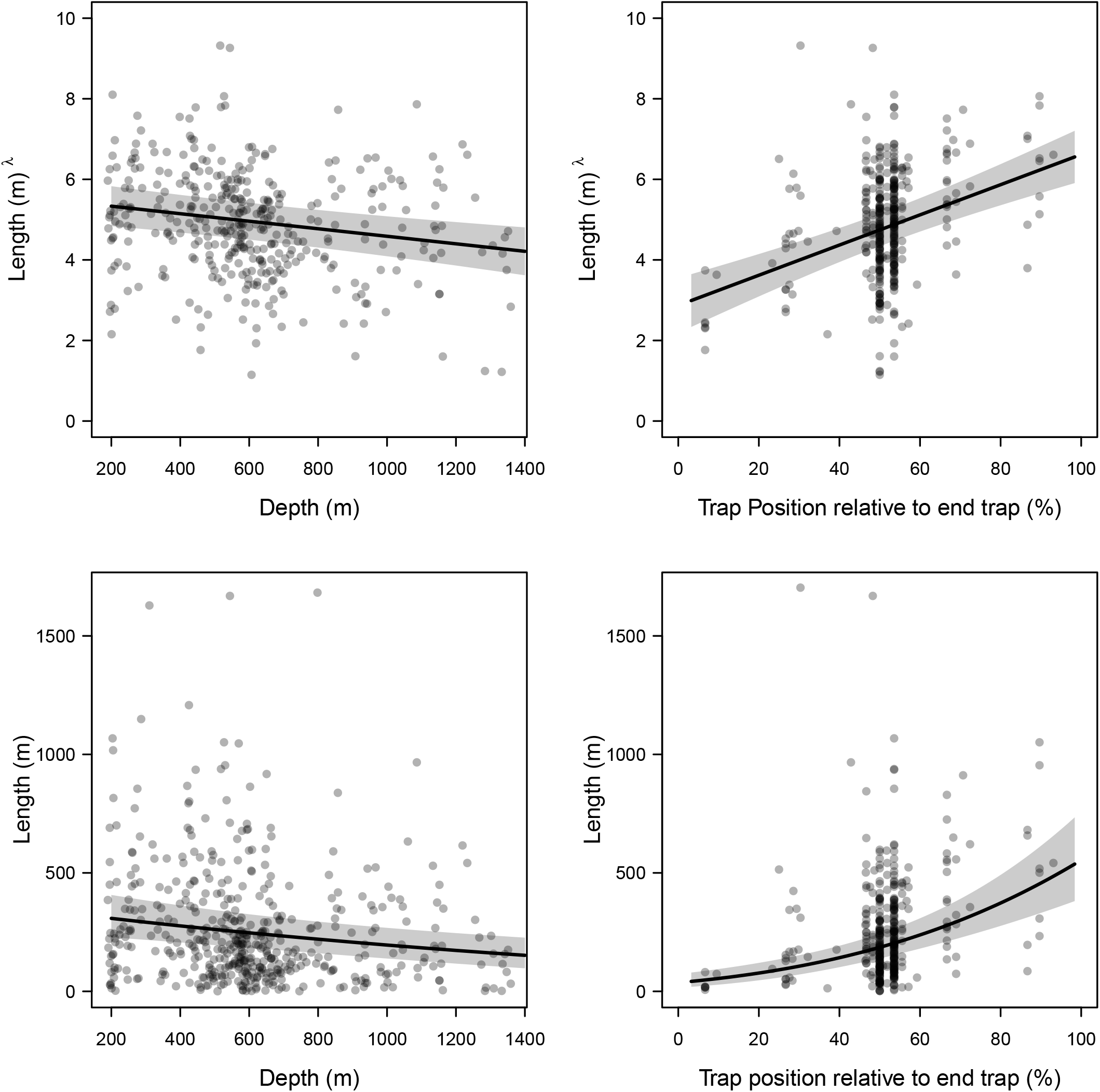
Plots on the left show estimated depth effect for the average trap position effect for a 60-trap set. Plots on the right show estimated trap position effect for the average depth effect from 200 to 1400 meters. Top plots show model fits to Tukey transformed (lambda = 0.3) drag length estimates, while bottom plots show non-transformed data. Black lines indicate the mean and grey polygons indicate 95% CIs (+/-1.96SE).

### 3.3. Footprint estimates

The estimated Sablefish trap footprint width for the different taxonomic groups range from 67 cm - 113 cm (Table 2) with a weighted footprint width of 97.6 cm. Most taxa had trap contact widths of approximately 1 m, with the exception of Hydrocorals that have shorter height distributions than corals, sponges, or sea whips.

We estimate that the combined footprints for commercial Sablefish longline trap and longline hook fishing gear contacted only 8.2% (95% CI: 4.4-11.5) of the seafloor area within grid cells fished by at least 3 unique vessels from commercial Sablefish fisheries from 2007-2023 and only 1.1% (95% CI:0.3-2.0) of the seafloor area was contacted more than once (Tables 3-4, Fig. 7). This accounts for 96% of the Sablefish longline and trap sets deployed from 2007-2023, after removing 4% of sets in sparsely fished areas with less than three unique fishing vessels ccording to DFO’s ‘rule of three’ for publicly released fisheries data (Tomasic, 2023). There were 604 (59%) grid cells that had between 0-5% of their area contacted by fishing gear at least once. In contrast, there were only 11 grid cells that had more than 50% (1.1%) of their area contacted. These estimates are based on the overlap method in eqn. (10), which is a more realistic approach to estimating the footprint; however, our estimates using the two methods (overlap vs. no overlap) do not differ much (Fig. 8) since the density of Sablefish fishing sets within most 4×4 km grid cells is low.

**Table 3:** The frequency of gear contact within Sablefish fishing areas for longline trap or hook gear from 2007-2023. Calculations use the random overlap approach with a total fished area of 16,336 km^2^, which is composed of 1021 4×4 km grid cells with Sablefish fishing from at least 3 unique vessels from 2007-2023.

| Number of times contacted | Area (km<sup>2</sup>) | % of fished area |
| --- | --- | --- |
| zero | 15008 (14453 - 15623) | 91.9 (88.5 - 95.6) |
| once | 1153 (662 - 1535) | 7.1 (4.1 - 9.4) |
| twice | 148 (47 - 273) | 0.9 (0.3 - 1.7) |
| at least three times | 27 (4 - 74) | 0.2 (0 - 0.5) |

**Table 4:**
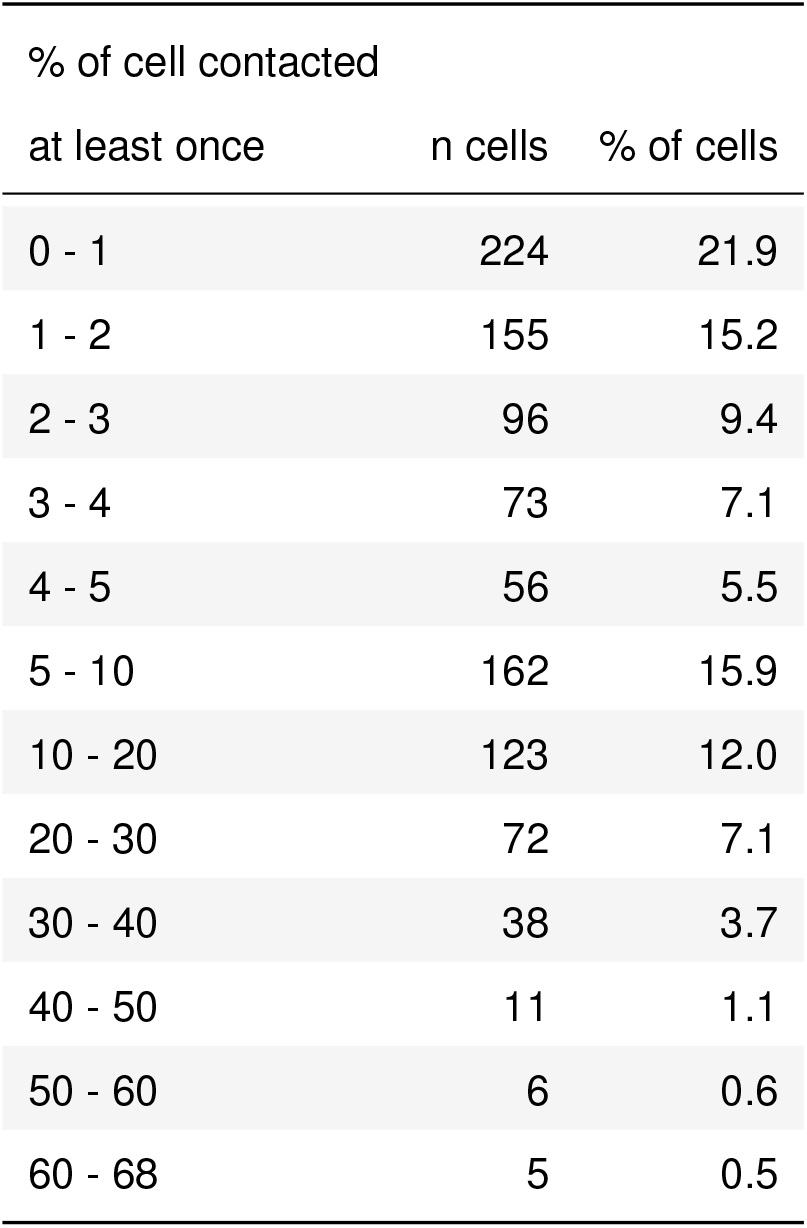
Summary of bottom contact estimates, assuming random overlap, for 4×4 km grid cells fished by Sablefish longline trap and hook gear from 2007-2023. Note that this table excludes grid cells with Sablefish fishing from less than 3 unique vessels from 2007-2023.

**Figure 6:**
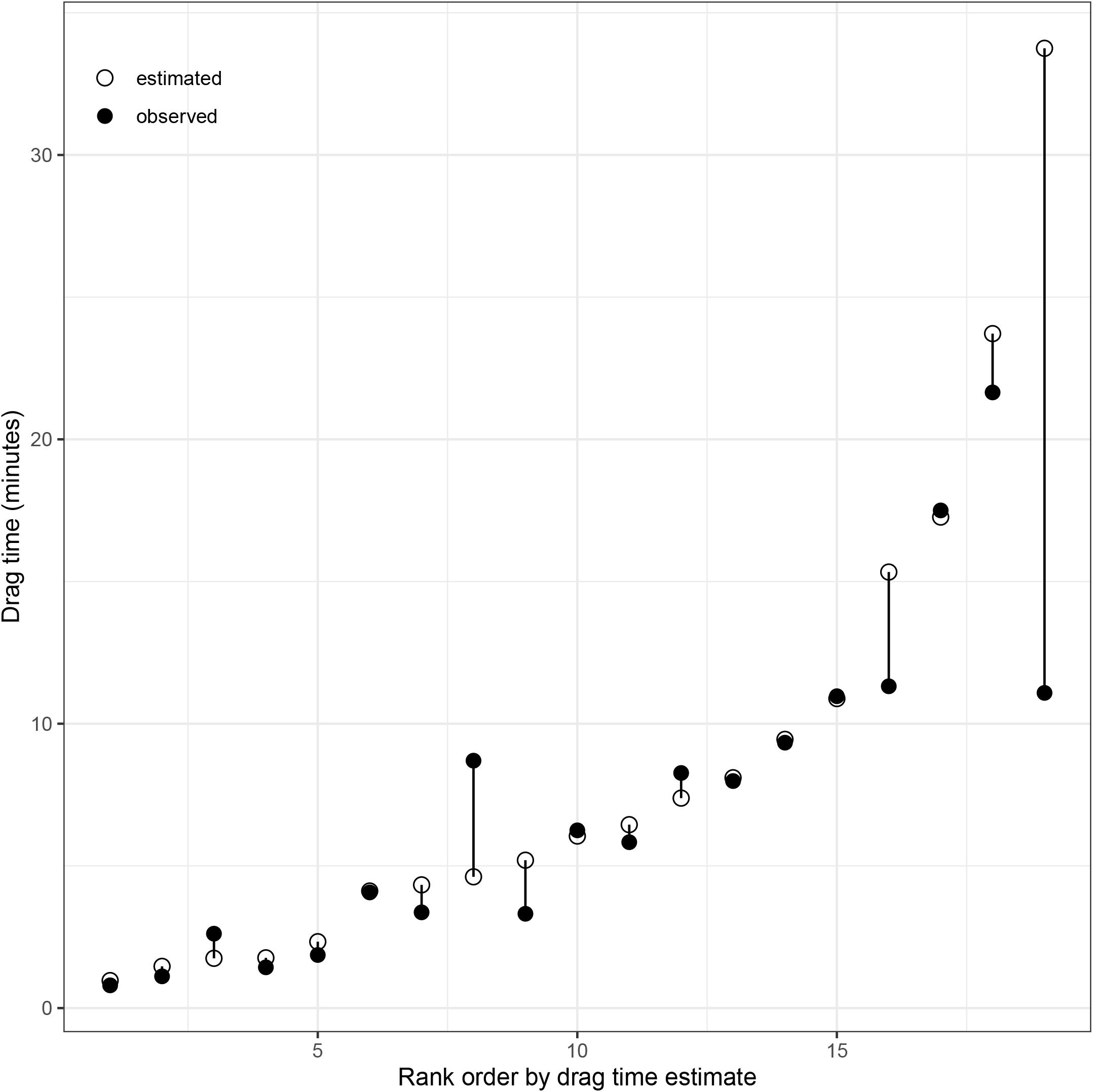
Differences between the estimated drag times from movement classification algorithm and observed drag times from video for 19 traps deployed in 2021-2022. Values are ordered by the estimated drag time from lowest to highest.

**Figure 7:**
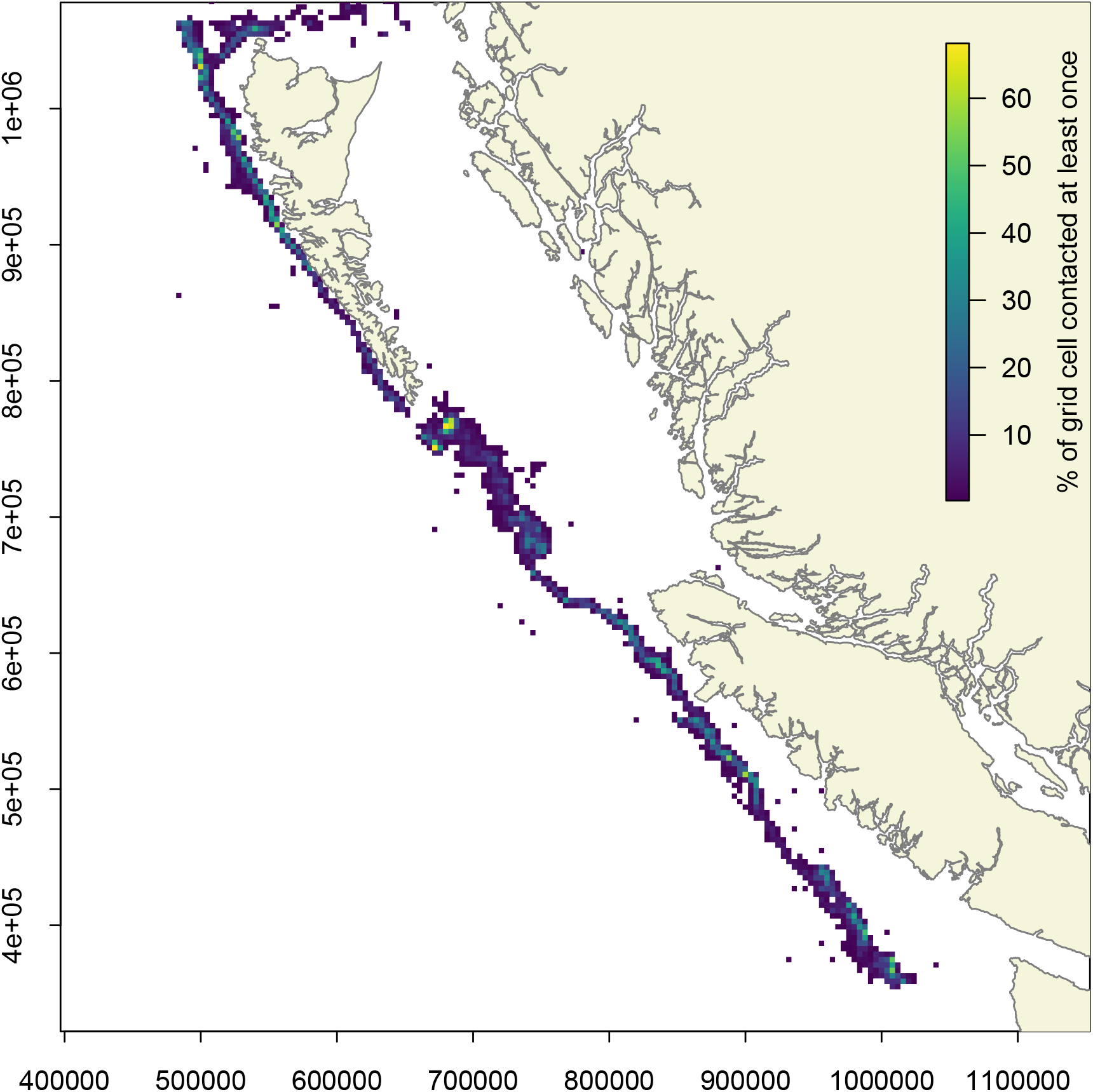
Proportion of 4 × 4 km grid cells contacted by Sablefish longline trap or hook gear at least once from 2007-2023. Note that this excludes 4% of the total fishery sets, which occur in grid locations fished by fewer than 3 vessels. Coastline data are from the nepacLL dataset (Wessel and Smith 1996) in the PBSmapping R package (Schnute et al. 2003). Map projection is NAD83 BC Albers (EPSG:3005).

**Figure 8:**
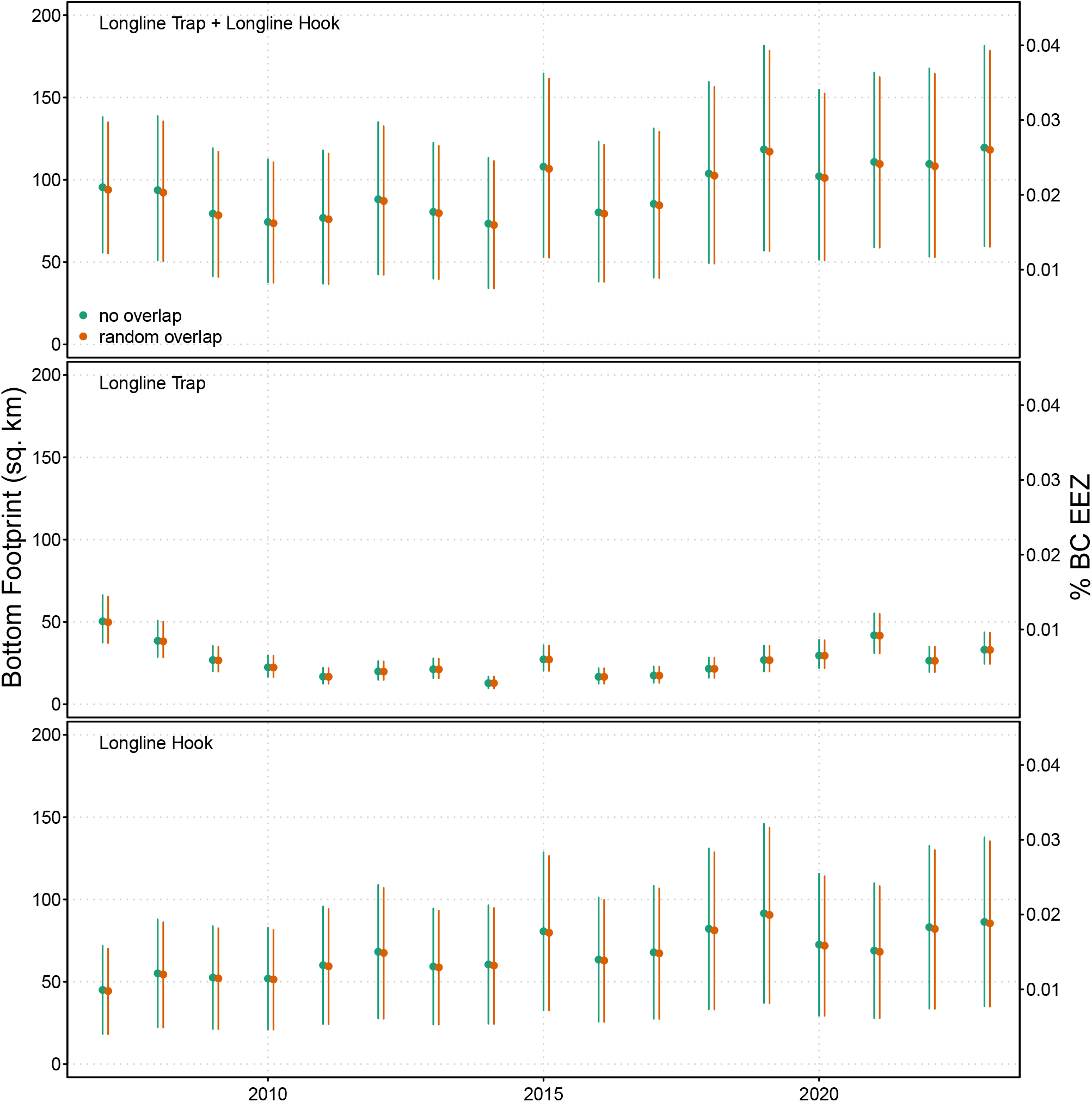
Mean estimates of annual footprints with 95% CI from Sablefish longline fishing gear for hook and trap combined (top), trap only (middle), and hook only (bottom) using the no overlap and random overlap methods calculated for 4 × 4 km grids. Proportion of bottom contact relative to BC Exclusive Economic Zone (EEZ) shown on right y-axes are based on EEZ area of 454,180 km^2^.

Mean estimates of annual bottom contact from the BC Sablefish fishery range from 73 km^2^ (95% CI: 34-112) to 118 km^2^ (95% CI: 59-180) for the 2007-2023 period, occuring in less than 0.03% of BC Waters (Fig. 8) and less than 0.6% of the Sablefish survey area (21,480 km^2) that covers depths between 183-1372 m on British Columbia’s continental shelf (Wyeth et al., 2007).

## 4. Discussion

Our bottom contact estimates indicate that Sablefish fishing risks to bottom habitat are low in the majority of fishing areas, since 91.8% of the area fished by at least 3 vessels within the BC Sablefish fishery is expected to have had zero bottom contact over the last 17 years. Furthermore, within the fished areas that are contacted by fishing gear (*k* >= 1), the majority are only contacted once. This indicates that most habitats contacted by Sablefish gear can be expected to have a minimum of 17 years to recover between subsequent bottom contact events, allowing longer periods for recovery. Considering this recovery time, there is potentially lower risk for long-term fishing impacts to bottom habitats in these areas (Kaiser et al., 2006a; Rooper et al., 2011; Kaiser et al., 2018).

Other fishing areas may have greater risk from fishing impacts, either because they are contacted more frequently by fishing gear or because they contain longer-lived species, such as some deep-sea corals that may require longer recovery times of 20-40 years (Rooper et al., 2011; Kaiser et al., 2018; Baco et al., 2019). For example, this could apply to the small percentage (1.1%) of Sablefish fishing areas that are contacted twice or more over 17 years. In these cases, a quantitative risk assessment (Welsford et al., 2014) approach could be used to provide a greater understanding of fisheries impacts and inform management decisions. Ideally, this should include i) estimates of bottom contact within fishing areas, ii) detailed mapping of the density and distribution of sensitive benthic habitats (Rooper et al., 2017b; Rowden et al., 2017b; Doherty et al., 2021), iii) estimates of mortality from gear contact (Martin-Smith and Welsford, 2014), iv) estimates of recovery following disturbance (Rooper et al., 2011), and v) an assessment of current and future population status relative to some reference level (e.g., unfished biomass) for different fisheries management strategies. Here we have demonstrated an approach for i), which could be applied for any type of longline fishery, while ii-v) will be addressed in future phases of the CSA bottom impacts research program. While quantitative risk assessments for SBHs involve multiple steps that might take place over several years or multiple phases of research, new information is generated at each step (i-v) that can progressively improve management decisions for habitat conservation.

Our study demonstrates an approach for estimating fixed gear bottom contact that can be widely implemented across longline fisheries. This serves as an essential initial step for habitat risk assessment, by providing fine-scale spatial information on fishery impacts to inform management decisions. For instance, there are only 11 of the 4 × 4 km^2^ grid cells fished where at least half of their area was contacted by Sablefish fishing gear from 2007 to 2023. This finding highlights potential higher risk areas that may warrant improved data collection and more detailed risk assessment. In such heavily impacted areas, information on habitat distribution and density (Rooper et al., 2017a; Rowden et al., 2017a; Doherty et al., 2021) and their overlap with bottom contact areas are essential next steps for effective risk assessment.

The steps for habitat risk assessment can be approached iteratively, allowing for the rapid production of an initial risk assessment focused on fisheries footprints, without the need to undertake all steps during early iterations of risk assessment. These initial risks assessments can provide valuable information for management decisions before considering other factors such as habitat distribution, mortality during gear contact, and potential recovery time, which are often data limited. As demonstrated for the BC Sablefish fishery, the initial step of estimating bottom contact area provides fine-scale information on potential habitat risks in fishing areas. Estimating bottom footprints for longline gear requires an approach for estimating gear contact per fishing event and spatial fishing effort data, the latter of which is readily available for many fisheries. For fisheries lacking specific information on gear bottom contact, scenarios for longline footprints could be developed based on our approach for estimating Sablefish longline trap footprints or other studies estimating longline hook footprints (Ewing et al., 2014). For instance, in the absence of movement data for Sablefish longline hook gear in BC, our footprint estimates used lateral movement estimates from longline fisheries for Patagonian Toothfish (Ewing et al., 2014). Future phases of research could then focus on collecting data needed to provide more robust risk assessment, such as deploying sensors and video on fishing gear to collect fisheries-specific data on gear bottom contact and presence-absence data for habitat mapping.

Footprint estimates for bottom longline fisheries in British Columbia could be improved in three ways. First, in this study we used estimates of lateral line movement (e.g., footprint width) for longline hook gear in the Patagonian Toothfish fishery; however, specific estimates for BC fisheries could be developed in the future by deploying cameras and other motion-sensing equipment on longline hook gear. Second, our analyses focussed on Sablefish and Pacific Halibut/Sablefish combined fishing trips using longline hook and trap gear. Larger footprints might occur in some areas where there are other longline (Pacific Halibut, Rockfish), trap (e.g., prawn), or trawl fishing that overlap with the Sablefish fishery. This is more likely an issue in shallower fishing grounds less than 700 m, as other BC fisheries typically do not fish at deeper depths down to 1400 m where Sablefish occur. Finally, future phases of the CSA bottom impacts research program will focus on improved camera and sensor systems, potentially allowing improved estimates of bottom contact and more data by facilitating wider adoption for both research and fishery application.

There is potential to improve the utility and accuracy of the movement classification algorithm by using a supervised machine learning approach on the outputs from the existing movement classification algorithm and observations from video footage. This might allow for an approach that uses only accelerometer data to estimate movement, without the need for video data which is time consuming to review and process. Additionally, a classification algorithm that combines accelerometer data with other low cost sensors such as gyroscopes or magnetometers to detect movement and orientation might provide more reliable movement classification and estimates of bottom contact (López Revuelta, 2017; Madgwick et al., 2011) that reduce misclassification errors, such as the outlier error from the 2021 Sablefish survey detected in the current study. Accelerometers, gyroscopes, and magnetometers are both cost-effective ($10-100) and compact, offering a more practical solution for large-scale deployment on fishing gear compared to depth sensors and video camera systems. Future applications might include the installation of low cost sensors on multiple traps along a groundline, providing real-time estimates of gear movement for each trap on the string.

## 5. Conclusion

We found that the BC Sablefish fishery bottom footprint occupies a very small proportion of total BC waters and that fishing risks are low within most of the areas fished. We demonstrate an approach for collecting data and estimating fishing gear bottom contact that can be widely applied to other longline trap and hook gear fisheries. Our findings address key data gaps in bottom impacts research for fixed gear fisheries, allowing fishing risks to be quantified over fine spatial scales. Such quantitative approaches for risk assessment can provide essential information for management decisions aimed at determining acceptable trade-offs between preservation of SBH and fishery benefits, which may allow fisheries to maintain access to fishing grounds where they can demonstrate that habitat risks are low.

## 6. Acknowledgements

We thank Wild Canadian Sablefish Ltd. (WCS) for their financial and in-kind support at all stages of this project from the initial design to field deployments and analysis. We especially thank the fishing masters who expertly deployed and retrieved camera traps on commercial fishing sets. Additional funding was provided by the British Columbia Salmon Restoration and Innovation Fund (BCSRIF). This project was also made possible by Kristina Castle and Malcolm Wyeth, who assisted with camera design, data collection and preparation, equipment preparation, and training of at-sea observers. We also thank Van Nguyen and Sijia Wang for processing sensor and video data, and for running the movement classification algorithm. We thank Kendra Holt and Chris Rooper for helpful comments that improved the manuscript.

## 7. Author contributions

Conceptualization: BD, AR, SPC Data curation: BD, LL Formal analysis: BD, LL, KA Funding acquisition: BD, AR Investigation: BD, AR, LL Methodology: BD, KA Project administration: BD, AR, SPC Software: BD, KA Supervision: BD, AR, SPC Visualization: BD, LL, KA Writing - original draft: BD Writing - review & editing: BD, AR, LL

## 9. Data availability statement

The CSA bottom contact research data (video, accelerometers, depth-sensor) used in the movement classification algorithm are not publicly available at this time. Researchers interested in using these data may contact the authors to discuss potential collaboration opportunities. The commercial fishery effort data for British Columbia groundfish fisheries logbooks are stored in the Fisheries and Oceans Canada (DFO) Fishery Operations System (FOS) database. Access to this data is governed by the Access to Information and Privacy Act and may be requested through DFO.

