## supplementary material S1 for "Treading lightly: Quantitative estimates of seafloor contact for longline trap and hook fishing gear"

### 1. Supplementary Material S1

This supplementary includes:

- maps of bottom contact by gear type for longline trap and longline hook gear
- estimates of areas contacted at least twice or three times for 4 x 4 km grids for longline trap and hook gear combined for 4x4 km

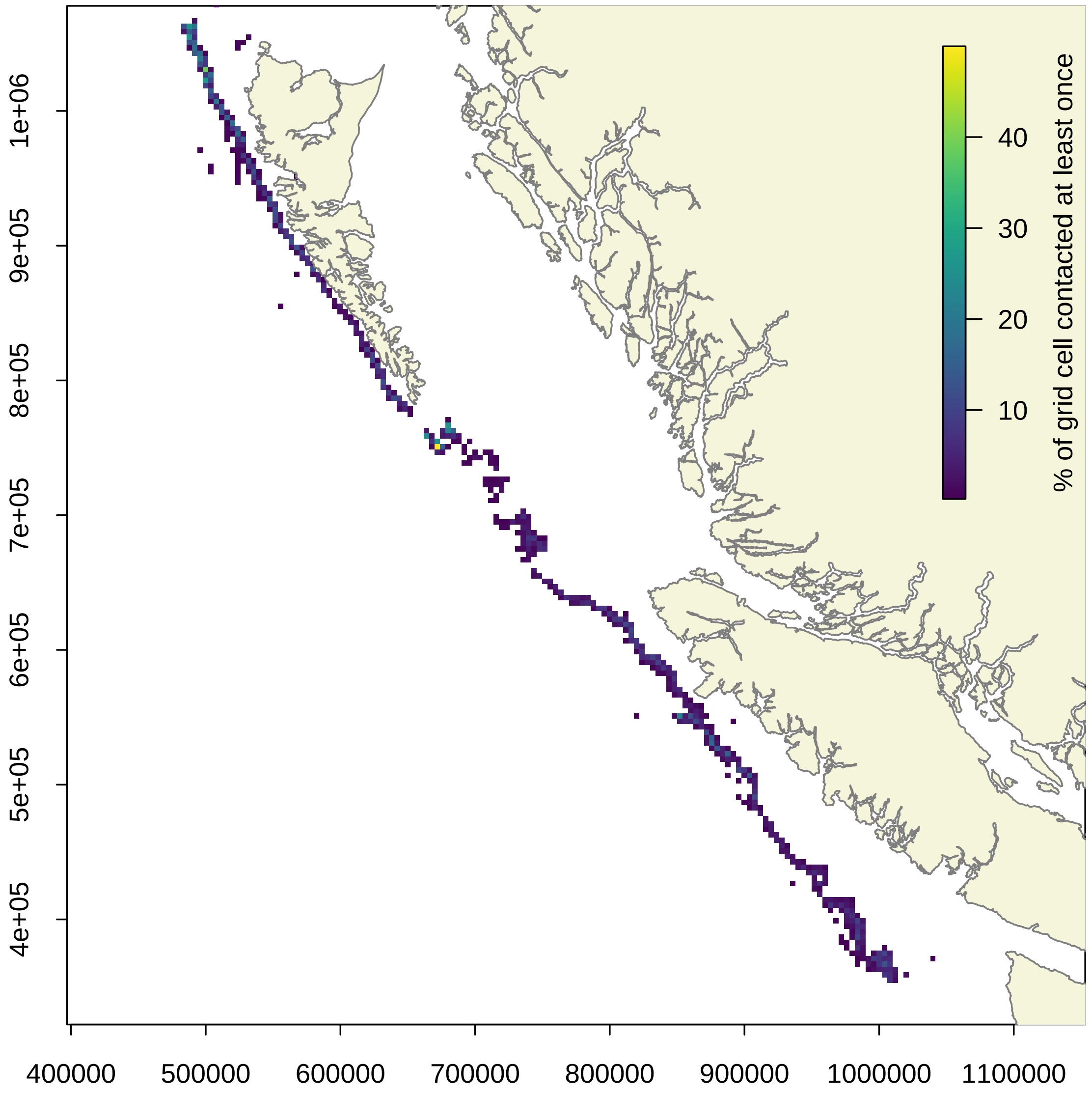


Figure 1: Proportion of 4 x 4 km grid cells contacted by Sablefish longline trap gear at least once from 2007-2023. Note that this excludes 7% of the total fishery sets, which occur in grid locations fished by fewer than 3 vessels. Coastline data are from the nepacLL dataset (Wessel and Smith 1996) in the PBSmapping R package (Schnute et al. 2003). Map projection is NAD83 BC Albers (EPSG:3005).

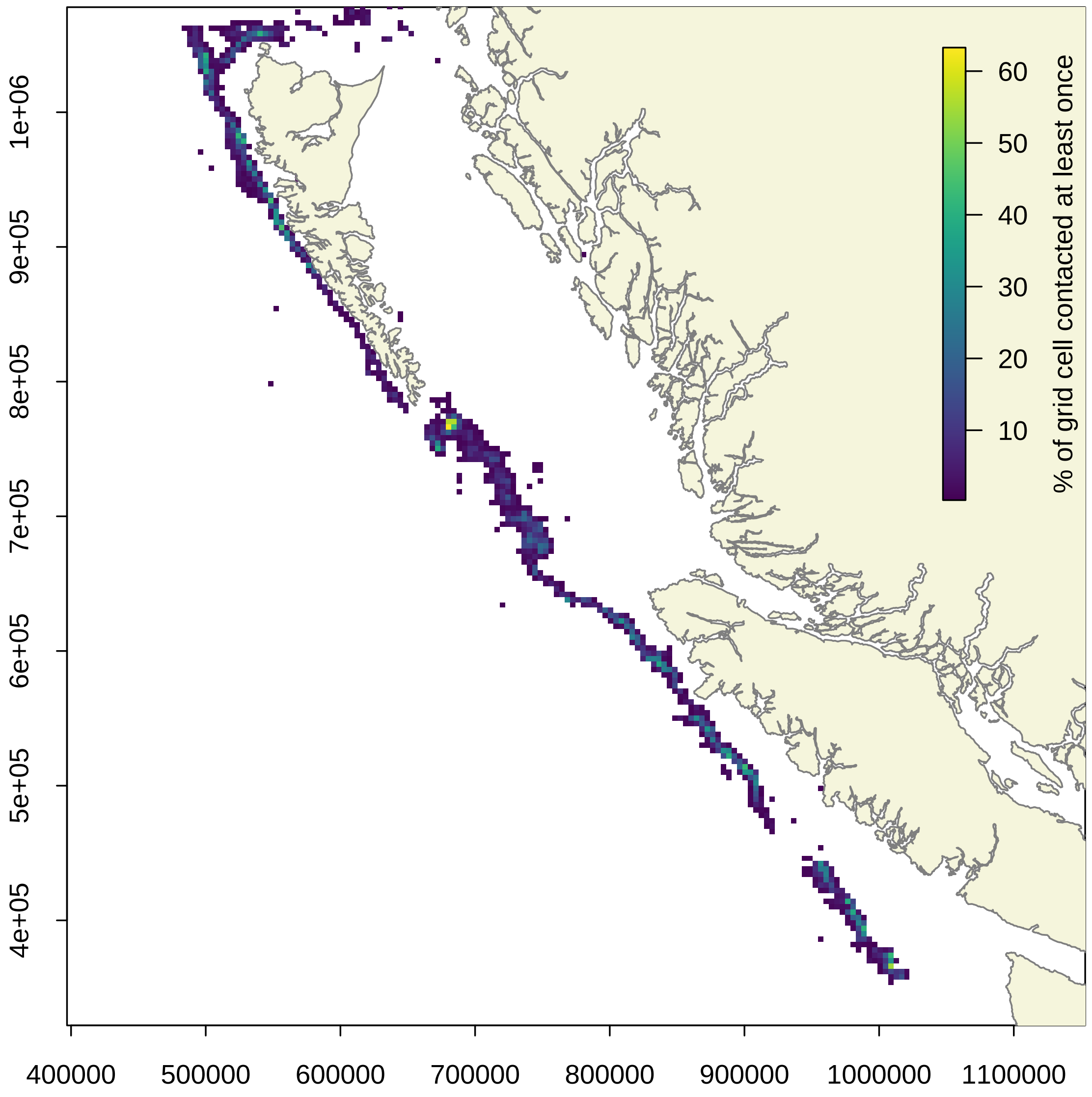


Figure 2: Proportion of 4 x 4 km grid cells contacted by Sablefish longline hook gear at least once from 2007-2023. Note that this excludes 5% of the total fishery sets, which occur in grid locations fished by fewer than 3 vessels. Coastline data are from the nepacLL dataset (Wessel and Smith 1996) in the PBSmapping R package (Schnute et al. 2003). Map projection is NAD83 BC Albers (EPSG:3005).

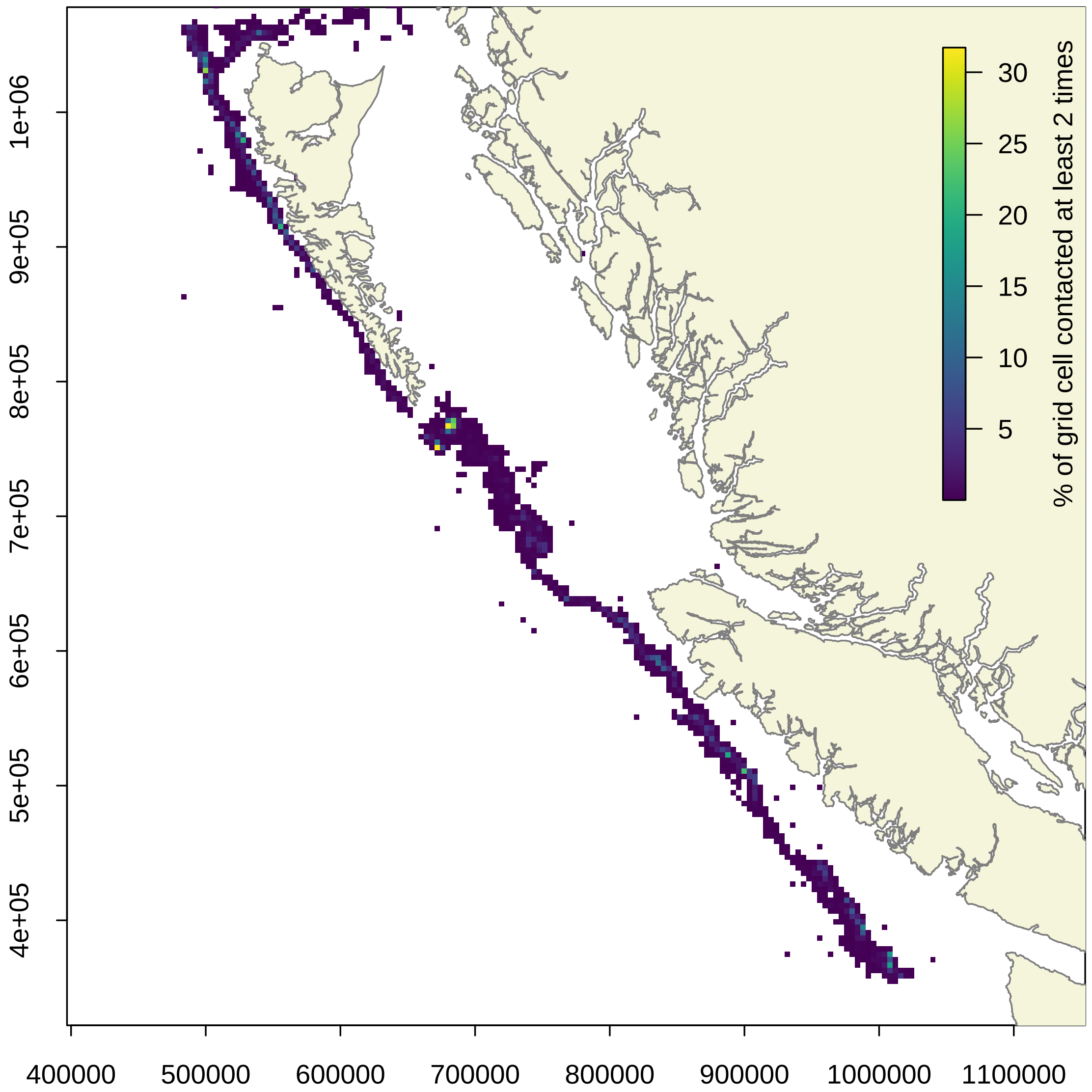


Figure 3: Proportion of 4 x 4 km grid cells contacted by longline Sablefish trap or hook gear at least two times from 2007-2023. Note that this excludes 4% of the total fishery sets, which occur in grid locations fished by fewer than 3 vessels. Coastline data are from the nepacLL dataset (Wessel and Smith 1996) in the PBSmapping R package (Schnute et al. 2003). Map projection is NAD83 BC Albers (EPSG:3005).

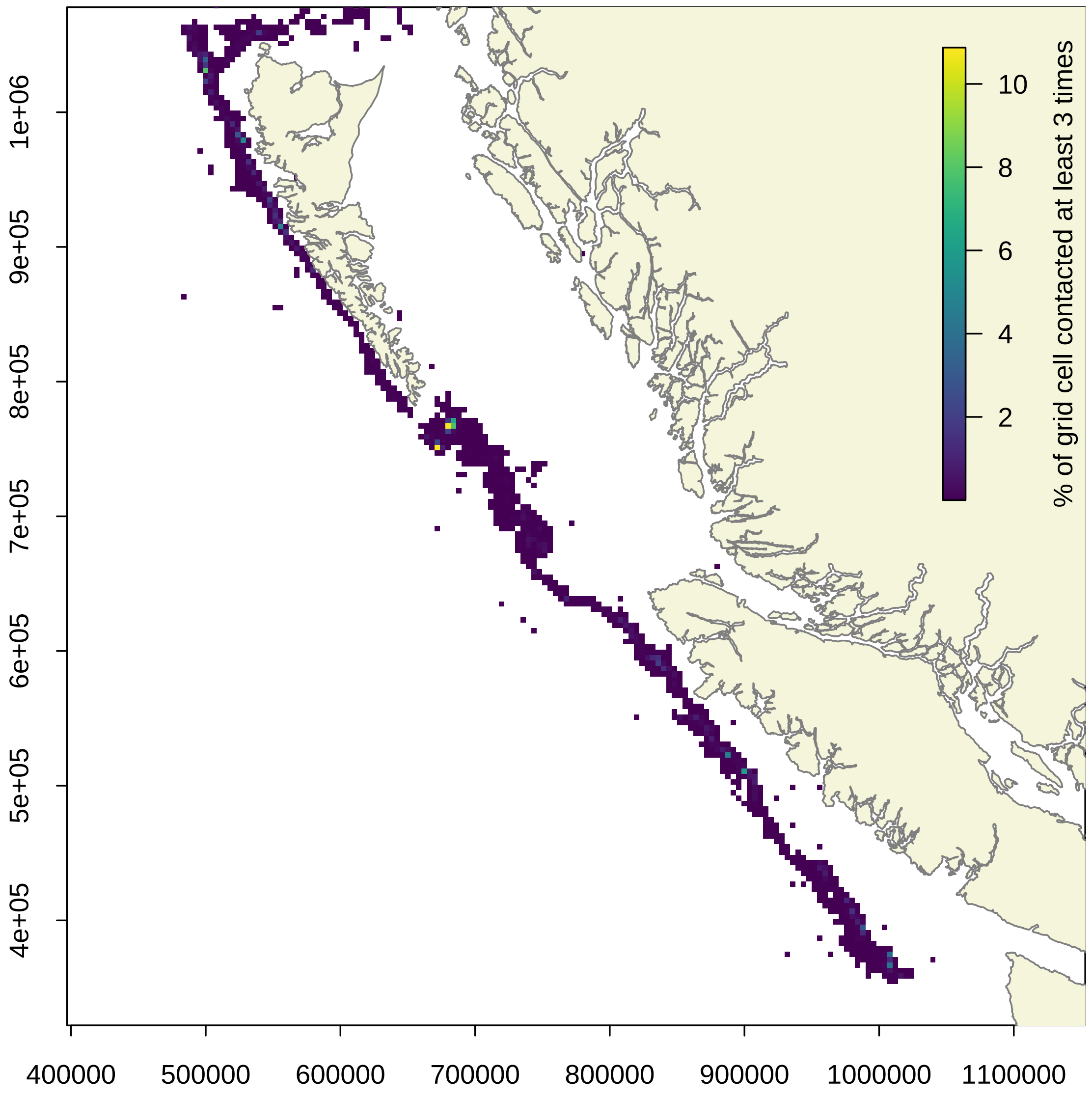


Figure 4: Proportion of 4 x 4 km grid cells contacted by longline Sablefish trap or hook gear at least three times from 2007-2023. Note that this excludes 4% of the total fishery sets, which occur in grid locations fished by fewer than 3 vessels. Coastline data are from the nepacLL dataset (Wessel and Smith 1996) in the PBSmapping R package (Schnute et al. 2003). Map projection is NAD83 BC Albers (EPSG:3005).
